# A role for ecologically-tuned chunking in the evolution of advanced cognition demonstrated by modelling the cleaner fish market problem

**DOI:** 10.1101/2021.05.30.446324

**Authors:** Yosef Prat, Redouan Bshary, Arnon Lotem

## Abstract

What makes cognition ‘advanced’ is an open and not precisely defined question. One perspective involves increasing the complexity of associative learning, from conditioning to learning sequences of events (‘chaining’) to representing various cue combinations as ‘chunks’. Here we develop a weighted-graph model to study the conditions for the evolution of chunking ability, based on the ecology of the cleaner fish *Labroides dimidiatus*. Cleaners must learn to serve visitor clients before resident clients, because a visitor leaves if not attended while a resident waits for service. This challenge has been captured in various versions of the ephemeral-reward task, which has been proven difficult for a range of cognitively capable species. We show that chaining is the minimal requirement for solving the laboratory task, that involves repeated simultaneous exposure to an ephemeral and permanent food source. Adding ephemeral-ephemeral and permanent-permanent combinations, as cleaners face in the wild, requires individuals to have chunking abilities to solve the task. Importantly, chunking parameters need to be calibrated to ecological conditions in order to produce adaptive decisions. Thus, it is the fine tuning of this ability which may be the major target of selection during the evolution of advanced associative learning.

## Introduction

In an effort to understand the evolution of cognition, a wide range of studies has been focused on identifying cognitive abilities in animals that appear “advanced”, and exploring the ecological conditions that could possibly favour their evolution (e.g., [1–6]). Yet, mapping sophisticated cognitive abilities along phylogenetic trees and their relation to social or ecological conditions (e.g., [7,8]) does not explain how such abilities evolved to become “advanced” through incremental modifications of their mechanistic building blocks. Earlier views of cognitive evolution were based on some postulated, loosely defined genetic adaptations, such as language instinct [9,10], mind reading abilities [11], or mirror neurons [12,13], but those are increasingly replaced by approaches relying on explicit associative learning principles that can gradually form complex representations of statistically learned information [14–22]. In line with these recent views, in order to understand the critical steps in cognitive evolution, one should identify specific modifications that can elaborate simple learning processes and make them better in some way.

A relatively simple and well-understood example is the extension of simple conditioning through second-order conditioning in a process known as chaining [23,24]. In this process, a stimulus associated with a primary reinforcer (such as a sound associated with receiving food), becomes a reinforcer by itself, and then a stimulus reinforced by the new reinforcer may become a reinforcer, and so on, allowing to represent sequences of statistical dependencies. Such sequences could, in turn, facilitate navigation [25,26] or even social learning [27].

Further elaborations of associative learning that may allow to construct complex representations and to support advanced forms of statistical learning and decision-making are less well-understood. It has become clear, however, that a critical ability required for such cognitive advances is the ability to represent two or more data units as a *chunk* or a *configuration* that has a meaning that is different from (or independent of) the meaning of its components. This ability has appeared in the literature under different names, such as *configurational learning* [28,29], *chunking* [19,30,31], or *segmentation* [32], all of which are quite similar, and involve the learning of configurations, patterns, and hierarchical structures in time and space [33].

In its simple form, known as configurational learning, this ability allows to learn, for example, that the elements A and B are associated with positive reward while their configuration AB is not rewarded and should therefore be avoided (a task known as negative patterning [34]). Configurational learning of this type is contrasted with *elemental learning*, which is based on the behaviour expected from simple associative learning [35,36]. Research on configurational learning has been focused mainly on identifying the brain regions supporting this ability (e.g., [29,37–39]), giving relatively little attention to the cognitive processes generating configural representations (but see [29]). More attempts to consider these possible processes has been made in the context of chunking or segmentation (e.g. [14,32,40]), but only recently, theoretical work has started to address the question of how chunking mechanisms evolve under different ecological conditions, and what is their role in cognitive evolution [19,41,42].

A unique model system that may provide a remarkable opportunity to study the evolution of chunking is that of the bluestreak cleaner wrasse (*Labroides dimidiatus*) which feeds on ectoparasites removed from ‘client’ fish [43]. Field observations and laboratory experiments have shown that at least some of these cleaner fish are capable of solving a problem known as the market problem (or the ephemeral reward task) [44–47]. The market problem entails that if approached by two clients, cleaners must learn to serve a visitor client before a resident client, because the latter waits for service while the former leaves if not attended (see details in [44,46,48,49]). In the lab, clients of different types were replaced with plates of different colours, each offering one food item and made to act like either a visitor or a resident [45,50]. Interestingly, individuals captured in different habitats demonstrated different learning abilities of the market problem in the lab, and adult cleaners seem to learn better than juveniles [46,48,51,52]. Such intraspecific variation in cognitive abilities suggests some role for the ecological and the developmental circumstances in the fish life history.

The lab market task may first appear as a two-choice experiment, testing whether animals can learn to choose the option that yields the largest total amount of food. Nevertheless, while preferring a larger amount in a simple two-choice task seems almost trivial for most animals [53], the market version, in which a double amount is a product of a sequence of two actions (i.e. choosing a visitor and then approaching the resident) has been proven difficult for a range of species [52,54–56] (but see [57–59]). Follow-up studies on pigeons and rats (reviewed in [60]) showed that letting the subject make a first decision but delaying the consequences, i.e. delaying the access to the rewarding stimuli, strongly improves performance [60,61]. One interpretation of these results is that the delay helped animals to connect their initial choice to both consequences; the first, and then the second reward, both of which occurred within a short time span after the relatively long delay.

While delaying the consequences of the initial choice may be helpful under some conditions, recent theoretical work suggests that under natural conditions, basic associative learning is insufficient for solving the market task, which instead warrants some form of chunking ability [62]. The reason for that is that the commonly used laboratory task presents a relatively simple version of the problem compared to the natural situation. It only presents visitor-resident pairs, for which choosing the visitor first, always entails double rewards and choosing the resident first always entails a single reward. In nature, on the other hand, cleaners face also resident-resident as well as visitor-visitor pairs, and most often only a single client approaches. As a result, choosing visitor first may not always entail double reward (e.g., in visitor-visitor pairs the second visitor is likely to leave) and choosing resident first may not always result in a single reward (e.g., in resident-resident pairs the second resident is likely to stay). Indeed, the theoretical analysis carried out by Quiñones et al. [62] showed that for solving the natural market problem it is necessary to have distinct representations of all different types of client combinations (visitor (*v*) + resident (*r*), *r+r, v+v, r*, and *v*), which means the ability to represent chunks. Yet, the analyses did not explain how such representations are created, and to what extent ecology causes variation in the cleaners’ ability to create such representations.

Following Quiñones et al.’s demonstration that chunking is necessary for solving the natural market problem, here we use the cleaner fish example as a means to study the evolution of an explicit chunking mechanism and the conditions that favour its success. Thus, we investigate how the very same problem – choosing between two options where one yields the double amount of food – set into an increasingly complex ecology selects for the evolution of increasingly advanced associative learning abilities. Our model is based on a weighted directed graph of nodes and edges, which initially form a simple associative learning model, and can then be modified to become an extended credit (chaining-like) model, or a chunking model. This approach allows to compare between clearly defined learning mechanisms and to pinpoint the modifications responsible for a presumed evolutionary step that improves cognitive ability. We analyse the three learning models’ performance in three tasks: the basic quantitative choice task, the laboratory market task and the market task embedded in a sequence of varying configurations (‘natural market task’). For the latter, we explored to what extent different densities and frequencies of client types select for different tendencies to form chunks (a critical parameter in the model), and how such different tendencies may affect the cleaners’ ability to solve the market problem.

## The core model

### Internal representation

Our core model consists of a weighted directed graph *G = (N, E)*, with nodes *N*, edges *E*, and additionally edge-weights *W*, node-weights *U*, and node values *F* (Fig. 1A). The basic model includes three internal nodes representing three external cues (perceived states): *N = {V, R, X}*, where: *V* – serving (feeding on) a visitor-client, *R* – serving a resident-client, and *X* – absence of clients (empty arena). These are the three cues required to represent the market problem and are therefore available to the cleaner fish in our simulations (Fig. 2). Edge-weights are updated according to the sequential appearance of the cues, i.e., whenever *n*_*j*_appears after *n*_*i*_the weight of the edge *n*_*i*_→*n*_*j*_, i.e. *W*(*n*_*i*_, *n*_*j*_), increases (by one unit, in our simulations). Thus, edge-weights represent the associative strength between nodes experienced one after the other. Node weights and values are attached to the cleaner’s decisions (see decision-making below) according to their occurrence and association of their outcome with food, i.e., whenever node *n*_*i*_is chosen, the weight *U*(*n*_*i*_) increases (by one unit, in our simulations) and the value *F*(*n*_*i*_) increases by the amount of food reward provided (which unless otherwise specified, is assumed to be one unit per client if served successfully, and zero otherwise). The value of a node could be regarded as the strength of its association with food, which can also be represented as the weight of the edge between the node and a reinforcer food-node (the weights of green arrows in Fig. 1A). The weighted directed graph constitutes the cleaner’s internal representation of the market environment. The cleaner’s decisions regarding which clients to serve depend only upon this representation (Fig. 1A).

**Figure 1.**
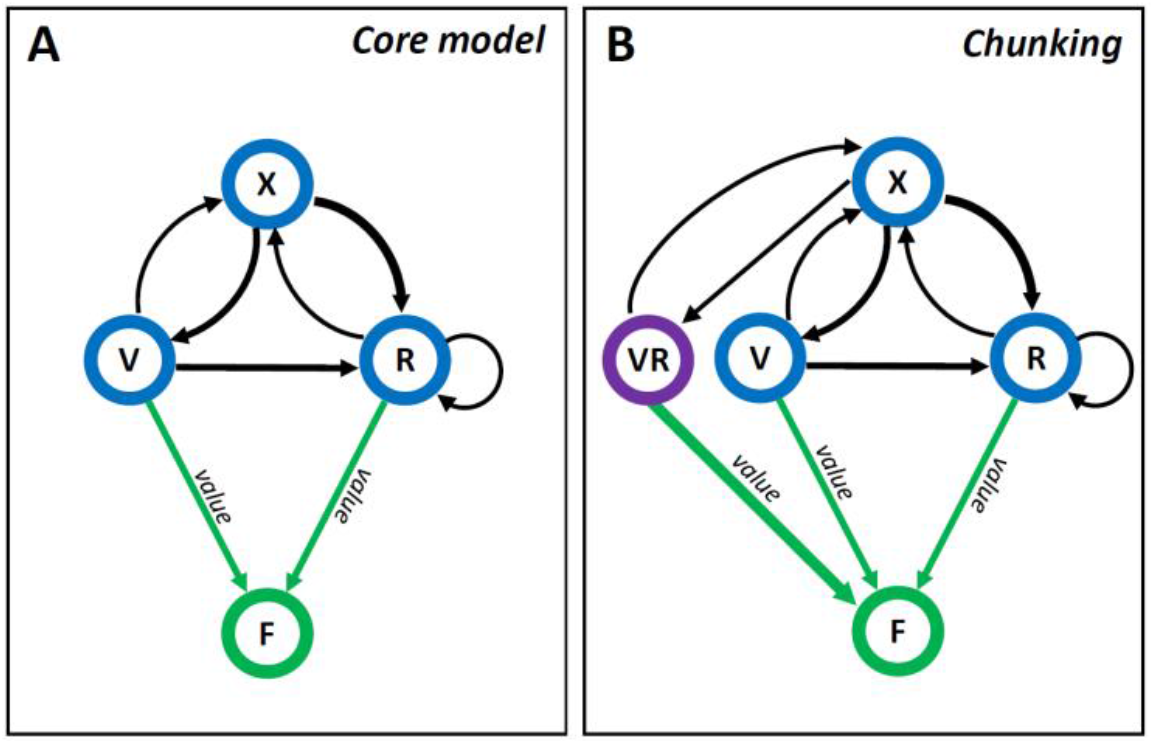
Model design - internal representation. **A**) The *core model* contains a network of three elements (blue circles) representing perceived states: *V* – serving a visitor-client, *R* – serving a resident-client, *X* – absence of clients. The value of each node is represented by the weight of its association (width of green arrows) with the reinforcer (food reward; green circle). Edge weights (width of black arrows) represent the strength of the associations between sequential states. This is also the internal representation of the *extended credit model*. **B**) An example of a possible representation in the *chunking model*: a new element (*VR*; purple circle) represents the configuration (the chunk) of ‘*V and then R’*.

**Figure 2.**
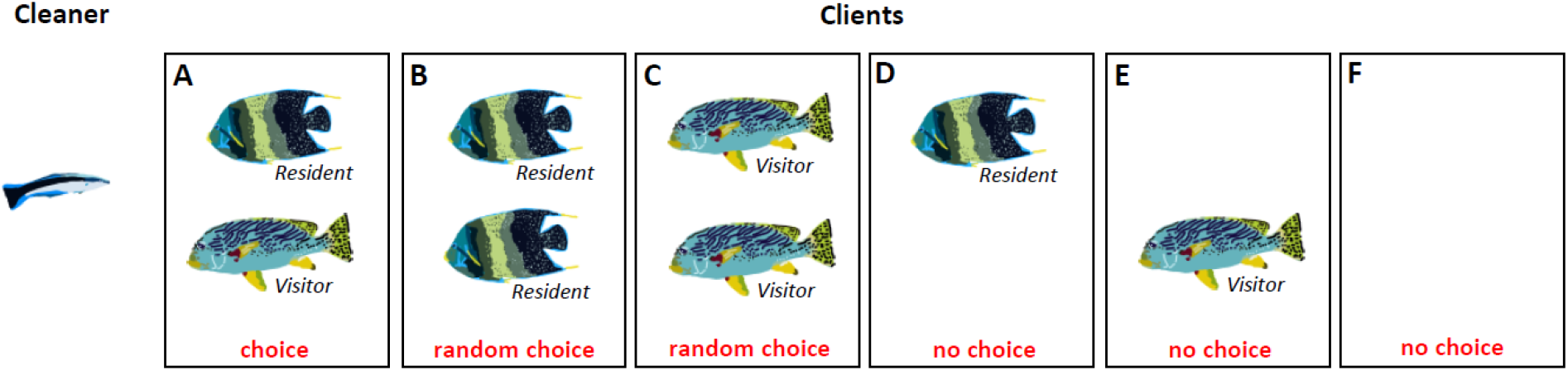
Model simulations. The cleaner in our simulations may encounter different combinations of client pairs awaiting its service: **A**) the cleaner must choose between two clients of different types according to the model’s decision process, **B and C**) The cleaner chooses with equal probabilities between two clients of the same type, **D and E**) the cleaner serves the only available client, and **F**) the cleaner waits for clients to visit its cleaning station.

Initially, the cues are considered unknown to the fish and their corresponding values, weights, and the weights of their connecting edges are set to zero. Most learning models use prior values for cues, which are commonly set to zero (often implicitly). Here, we model such a prior by imposing a threshold on the weight of a node before any increase in its value *F* can occur. Specifically, *F*(*n*_*k*_) is initialized to zero, and would not change as long as *U*(*n*_*k*_) < *Q*, i.e., at the first *Q* occurrences of *n*_*k*_. We set *Q* = 10 throughout all simulations, which implies that the value of a node will increase above zero only from the 11^th^ serving of a client.

### Decision making

When a cleaner fish is presented with two clients, it must choose which one to serve first. If both clients are of the same type (i.e., *v* (visitor) and *v*, or *r* (resident) and *r*) the cleaner chooses one with equal probabilities. However, when two contrasting types are present (i.e., *v* and *r*) the decision is made according to the values associated with serving each type, *F(R)* and *F*(*V*). A soft-max function is employed (see [62]) over the *normalized values f*(*n*_*i*_) and *f*(*n*_*j*_) such that the probability of choosing *n*_*i*_is:

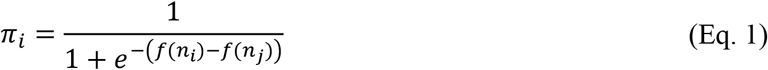

Where 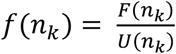 is the average payoff associated with the node *n*_*k*_. Note that the numerator, *F*(*n*_*k*_), is the sum of all obtained reward items associated with the cue *n*_*k*_ (i.e., the accumulated number of food items obtained after the cleaner has chosen *n*_*k*_), and the denominator is a count of all occurrences of *n*_*k*_ (i.e., the number of times the cleaner has chosen *n*_*k*_, regardless of whether this choice had been fulfilled).

The probability of choosing *n*_*j*_ is *π*_*j*_ = 1 − *π*_*i*_.

In the market problems presented here, both client types provide the same immediate reward. Thus, it is quite intuitive that learning only first order associations cannot provide any discrimination between them, and consequently, would fail in developing a preference for visitors (which is the essence of solving the market problem). Indeed, as we shall see in the results section, the core model was never successful in solving the market problem (either in its simple laboratory version or more complex natural setting). Yet, it serves as a null model and as a stepping-stone for the more advanced learning models.

### A linear operator model

To compare our core model with a similar known benchmark we used the linear-operator learning model [63], which is a basic and widely used learning model [64] that does not involve chaining or chunking. The learner updates the value *f(i)* of cue *i* at time *t* such that *f*(*i*)_*t*_ = (1 − *α*)*f*(*i*)_*t*−1_ + *αφ*(*i*)_*t*_, where *φ*(*i*)_*t*_ is the reward attached to cue *i* at time *t* and α is a learning-rate parameter. To choose between clients based on their updated values we used the same soft-max decision-making rule applied by the core model (see above).

## The extended credit model

A straightforward approach to consider higher order dependencies is to enable association of cues with their ‘future’ rewards. We call this model the ‘extended credit’ model. The network representation of the extended credit model is the same as that of the core model (Fig. 1A) but in this model the learner associates an obtained reward with the current cue as well as with the previous one. Specifically, while encountering a sequence (*n*_*i*_, *n*_*j*_), if *n*_*i*_ is rewarding then *F*(*n*_*i*_) increases, and if *n*_*j*_ is rewarding then both *F*(*n*_*i*_) and *F*(*n*_*j*_) increase (i.e. the credit assignment of the reward is extended also to the previous cue). Hence, if both cues are similarly rewarding the first one will be associated with double the food by the end of the sequence, as it was also associated with a delayed reward. Theoretically, credit assignment could be extended in more than one step backward and the credit could also change (e.g., decrease) with time (similarly to ‘chaining’ [65]). Note that although the model extends the credit to a previous cue, it does not represent, in the credit extension, the **identity** of the consecutive cue which donated the extra reward. Thus, the extended credit model cannot learn to distinguish between different sequences (sequential combinations or configurations) of cues (e.g., *V*→*R, V*→*X, R*→*R*, etc.). The decision-making process of the extended credit model is the same as in the core model (see above).

## The chunking model

Another way of identifying high order dependencies is via configurational learning, or chunking, as mentioned in the Introduction section. To model how acquired experience leads individuals to create chunks, we employ a chunking procedure in our model in which sequences occurring more often than expected, according to the distribution of their elements are ‘chunked’ into a new element (Fig. 1B). Specifically, a sequence (*n*_*i*_, *n*_*j*_) would become a new element ‘*n*_*i*_*n*_*j*_’ of the internally-represented network (i.e., a new node in the graph *G*) whenever:

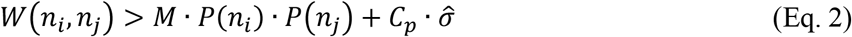

where *W*(*n*_*i*_, *n*_*j*_) is the number of observed occurrences of the sequence *n*_*i*_→*n*_*j*_, *M* is the total number of observed cues (or pair sequences), *P*(*n*_*k*_) is the observed frequency of the element *n*_*k*_, and 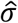 is the standard deviation of a binomial distribution, with the probability of an event *n*_*i*_→*n*_*j*_being *P*(*n*_*i*_)*P*(*n*_*j*_):

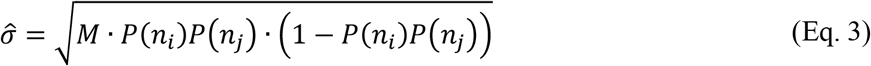

*C*_*p*_ *≥* 0 is a chunking avoidance parameter. This parameter is important, as it governs the behaviour of our model, or in other words, the conditions under which a chunk will be created. Note that when *C*_*p*_=0, any slight above chance co-occurrence of *n*_*i*_and *n*_*j*_would result in chunking. This is probably too much chunking because it can easily happen in nature for almost any two elements as a result of stochastic deviations from the frequency expected by chance. Using a *C*_*p*_ that is greater than zero implies that a chunk will be created only when the co-occurrence is higher than expected by a certain threshold.

Additionally, chunks would not be created as long as *W*(*n*_*i*_, *n*_*j*_) < *Q*, i.e., during the first *Q* occurrences of the sequence *n*_*i*_→*n*_*j*_. This rule enforces a minimal sample size before statistical inference could be done (for chunking).

In this model (see Fig. 1B), whenever a chunk is created it is treated as a new node and is being associated with food whenever chosen by the cleaner alongside food reward (but only after its first *Q* occurrences, as required for other elements). For instance, if the sequence *V* →*R* is chunked into a new element ‘*VR*’, further choices of the sequence *V* →*R* will increase the association of the element ‘*VR*’ with the reward by two units (as this is the observed reward during the processing of the sequence). On the other hand, if the sequence *R* →*V* is chunked into a new element ‘*RV*’ (which could happen in the *natural market problem*; see simulated environments below), further choices of the sequence *R* →*V* will usually increase the association of the element ‘*RV*’ with the reward by one unit only (since the visitor leaves if not served first).

The decision-making process of the chunking model is the same as in the core model (see above) but here, more choices may become available. For example, after the chunk ‘*VR*’ is created, a cleaner faced with a visitor and a resident client simultaneously can choose to serve the resident (*R*), or to perceive them as the chunk ‘*VR*’ and to execute the sequence *V* →*R* (i.e. approach the visitor and then the resident). On the other hand, if the chunk ‘*RV*’ was also created, an additional option exists, which is the choice of executing the sequence *R* → *V*. Importantly, in this case, soon after approaching the resident, the visitor would leave the arena so the outcome of choosing and attempting to execute the sequence *R* → *V* may end up with serving only *R* (depending on the simulated environment; see below) and being reward by only one unit (see above). We assume that if a chunk has already been created the cleaner never chooses the first element alone if presented with both elements (i.e., if ‘*VR*’ is already represented in the network, and ‘*RV*’ is not, the cleaner should only choose between ‘*R*’ and ‘*VR*’ when presented with both client types simultaneously).

## Simulated environments

Our simulations provide the cleaner fish with alternating clients awaiting its service (Fig. 2). The simulated arena includes two available spots, where each can be occupied by a visitor client, a resident client, or remain empty (simultaneous encounters with more than two clients are relatively rare in nature and are typically not addressed; see [44,45]). Each simulation consists of a sequence of discrete trials. On each trial, the arena is filled using a random sample according to the simulation specific setup (i.e., the probabilities of encountering each client type, as will be explained below). When the cleaner is presented with an empty arena (none of the two spots is occupied) it perceives the corresponding cue *X* (see Fig. 1A) and waits for the next trial (the next occupancy of the two spots). When the cleaner is presented with only a single client it immediately serves it, perceives the corresponding cue (i.e., *V* for choosing to serve a visitor, or *R* for choosing to serve a resident), experience the associated reward of serving it, and waits for the next trial. When the cleaner is presented with two clients, it chooses one according to the decision rule of the model (see above) if the clients are of different types (a visitor and a resident), or at random (with equal probabilities) if they are of the same type. The cleaner then serves the chosen one, and perceives the corresponding cue (i.e., *V* or *R*) and its associated reward. If the second client (not chosen) is a visitor it leaves and the cleaner waits for the next trial, but if the second client is a resident, the cleaner serves it as well, perceives the corresponding cue (*R*) and its associated reward, and waits for the next trial. Recall that whenever the cleaner chooses to serve a client and perceives its associated food reward, it adds one unit to the value *F* of the corresponding cue (e.g., to *F(V), F(R), F(VR)*, etc.).

We have simulated three different environments: *i*) A laboratory environment with a ‘basic two-choice task’, where a cleaner has to choose between two clients offering a reward of 1 and 2 units respectively, and no further approach to clients is allowed after this initial choice within a trial (only in this simulation, both client types are ephemeral). This two-choice task is expected to be solved by all types of leaners (i.e., preferring the client offering the double amount of food), thus serving as a ground-level associative learning test. *ii*) A laboratory environment with a ‘laboratory market problem’, where the cleaner faces a resident and a visitor client together (Fig. 2A), in each feeding trial, and after it finishes feeding it faces a single trial of empty arena (Fig. 2F; i.e., perceives the cue *X*). *iii*) A natural setting, henceforth termed ‘the natural market problem’, in which the cleaner may face all possible combinations (Fig. 2A-F): a visitor and a resident, two residents, two visitors, a single client (resident or visitor), and no clients. In addition, in the natural setting the cleaner does not necessarily have to wait between trials. This environment simulates more faithfully the situation in the wild, where each of the two spots is filled using an independent random sample, with a probability *P*_*V*_ for a visitor, a probability *P*_*R*_ for a resident, and a probability *P*_*0*_ for an empty spot (*P*_*V*_ + *P*_*R*_ + *P*_0_ = 1). When examining *the natural market problem*, we consider the distribution of the different client types and their combinations as resulting from two ecological parameters: the (relative) visitor frequency, 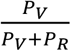 (the fraction of visitors out of all clients), and the overall client density, 1 − *P*_0_ (ranging from zero – when there are no clients and the cleaner always faces an empty arena, to one – the arena is always full).

## Results

We examined how the four learning models fare in the three simulated tasks: the basic two-choice task, the laboratory market problem, and the natural market problem.

### The basic two-choice task

All learning models solved successfully the basic two-choice task, as expected, exhibiting clear preference for the client offering double amount of reward, and showing virtually no differences in speed and accuracy of learning (Fig. 3A). In this task, there are no sequences of rewarding cues (as only the chosen client is consumed and the other leaves) thus the advanced models are practically reduced to the core model. Thus, the extended credit and the chunking model confer no extra benefit when facing a basic two-choice test between options that differ in the amount of reward.

**Figure 3.**
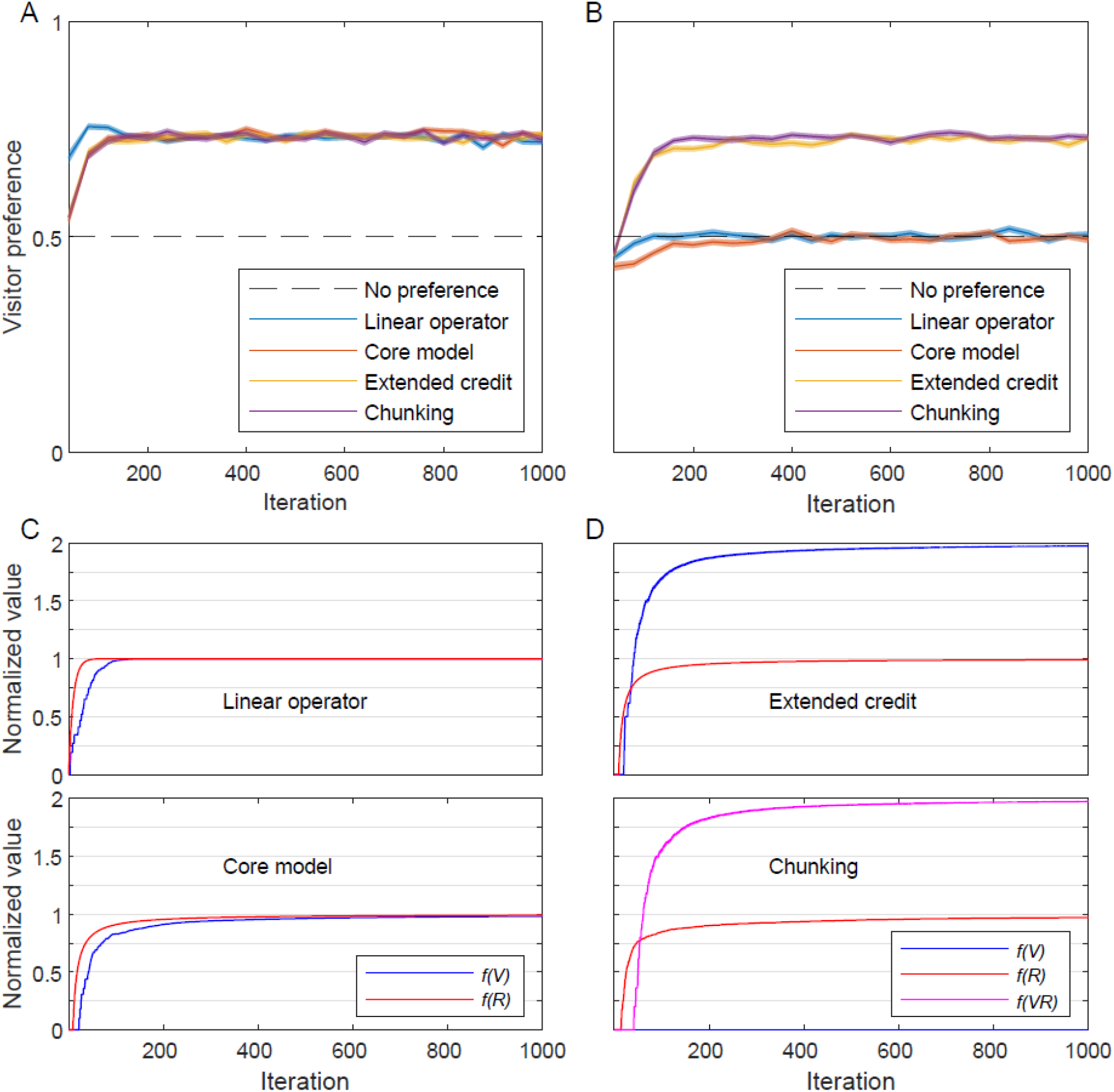
Simulating laboratory environments. Four types of learners are compared in the *basic two-choice task* **(A)** and the *laboratory market problem* **(B)**: blue – A linear operator learner (α=0.1; see text); orange – the core model; yellow – the extended credit model; purple – the chunking model (with *C*_*p*_=2); black dashed-line – the expected choices with no preference (0.5). The preference towards a visitor client, measured as the proportion of choosing a visitor out of all visitor-resident encounters, is plotted as a function of time (iterations), in bins of 40 trials. Both laboratory environments were simulated using 1000 feeding trials (with an empty trial after each feeding trial). The plots depict the mean of 100 simulations for each learner (shades – standard error of the mean). **C)** The value of the different cues as perceived by the non-successful models, the linear operator (top) and the core model (bottom), in a single simulation of the *laboratory market problem*: blue – *V*; red – *R*. **D)** The values perceived by the successful models, the extended-credit model (top) and the chunking model (bottom) in a single simulation of the *laboratory market problem*: blue – *V*; red – *R*; magenta – *VR*. Note that the chunking model, in this task, quickly creates the *VR* chunk, even before any value is attached to *V* itself.

### The laboratory market problem

Facing the laboratory market problem, the core model and the equivalent linear-operator learner did not develop a preference towards any of the given clients and thus failed to solve the problem (Fig. 3B, orange and blue lines, respectively). In contrast, both the extended-credit model and the chunking model were capable of solving the *laboratory market problem*, i.e., to develop a strong preference towards the visitor client (Fig. 3B, yellow and purple lines, respectively). The inability of the core and linear-operator models to solve the laboratory market problem is reflected in their indifferent *normalized value*s of *R, f*(*R*), and *V, f*(*V*), both of which approach 1 (Figure 3C). This result is expected since the value of each cue is updated independently of any past and future cue or reward, and both cues (client types) provide the exact same immediate reward. On the other hand, in the extended credit model that solves the problem successfully, the *normalized value* of *R, f*(*R*), approaches 1, while the *normalized value* of *V, f*(*V*), approaches 2 (Fig. 3D, top panel). This was made possible because serving a resident always provides a single food item in this setup (as the visitor leaves) while the credit of choosing a visitor is extended to the resident that waits to be served (thus crediting *V* with two food items). The success of the chunking model is based on a different process: it creates the chunk *VR* early and *f*(*VR*) quickly approaches 2 as the complete chunk provides two food items (Fig. 3D, bottom panel), pushing the preference towards a visitor client (the model choses the sequence *VR*, i.e., *V and then R*).

### The natural market problem

For the natural market problem, we only present the results of the learning models that were successful in solving the laboratory market problem (as expected, the core and the linear-operator models that failed to solve the laboratory problem also fail to solve the more complex natural problem, data not shown).

The extended-credit model that was sufficient for solving the laboratory market problem failed to solve the natural market problem (i.e., to prefer a visitor client) regardless of the overall client density or the relative frequencies of different clients (see examples in Fig. 4A, yellow lines). The reason for that is that in the *natural market problem* all pair sequences can occasionally appear, including a resident after a resident (and thus *R* is credited with 2 food items), a visitor after a visitor (and thus *V* is credited with 1), and even a resident and then a visitor (when there is no empty trial after serving the resident, which again credit *R* with 2). Thus, assigning credit for a cue for the value of the next cue causes the differences between *f(R)* and *f(V)* to vanish. Still, the sequence *V* →*R* may occur more often than the sequence *R* →*V* (at least as long as the cleaner do not prefer *R*), since whenever the two types of clients appear simultaneously, *V* →*R* occurs if the cleaner chooses to serve the visitor first, and *R* →*V* occurs only when a visitor appears by chance in a new trial after the cleaner has served a resident. As a result, *f(V)* might be slightly greater than *f(R)* in some situations. However, in order to respond to such slight differences, the model’s soft-max decision rule should be ‘hardened’ (become more similar to a maximum-based rule). This would suppress exploration and make the model always choose the most frequent client type (as its value increases faster), which is the resident in most cases since the visitor leaves if not served.

**Figure 4.**
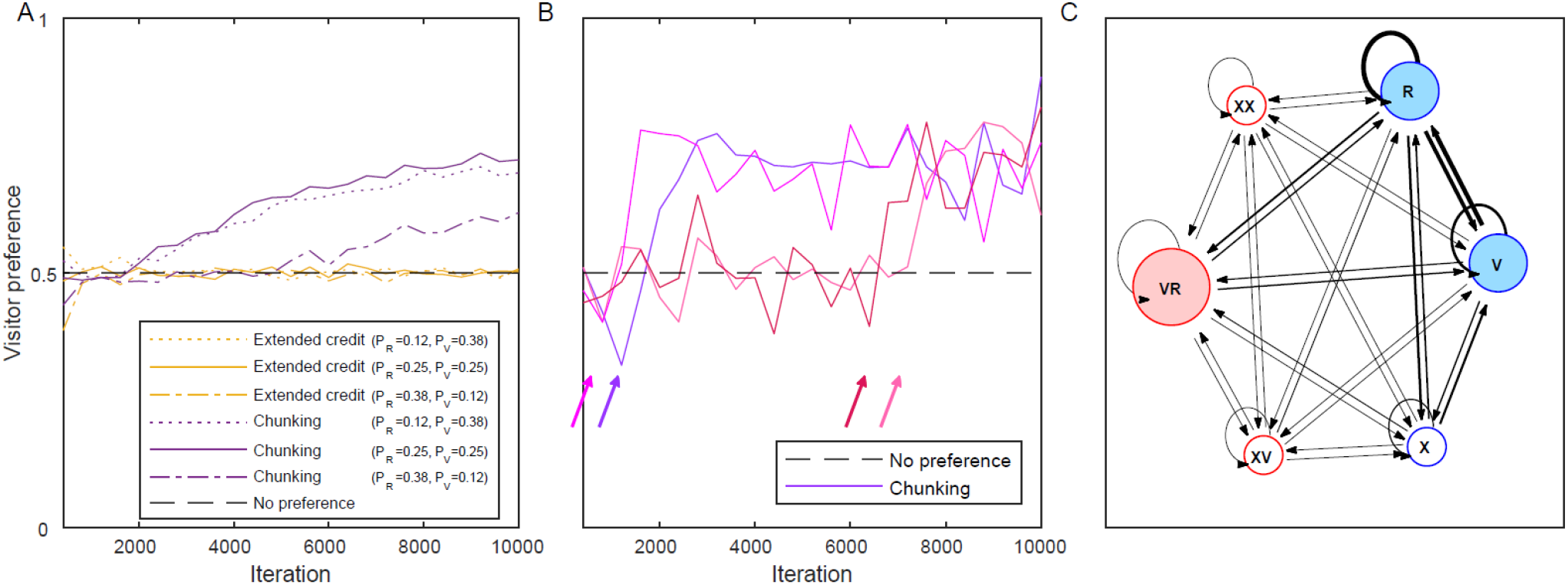
Simulations of the *natural market problem*. **A)** The preference for a visitor by the extended-credit model (yellow) and the chunking model (purple) are presented for client density (*1-P*_*0*_) of 0.5 and for three different distributions of client types: *P*_*R*_=0.12 and *P*_*V*_=0.38 (dotted lines), *P*_*R*_=0.25 and *P*_*V*_=0.25 (solid lines), *P*_*R*_=0.38 and *P*_*V*_=0.12 (dotted-dashed lines). Black dashed line – no preference (0.5). *C*_*p*_=2 (for the chunking model). **B)** Four simulations of the chunking model in the *natural market problem* (with *P*_*R*_=0.25 and *P*_*V*_=0.25). Note how the preference towards a visitor sharply increases after the creation of the *VR* chunk (depicted with an arrow of a corresponding colour for each simulation). **C)** The internal representation of the chunking model at the end of a simulation as in **(B)**. Blue – basic (initial) elements, red – chunk elements, filled nodes – the relevant elements for the decision process. The size of the circle is relative to the value (association with food reward) of the element. The width of the directed edges (black arrows) is relative to the weight (*W*) of the transitions between elements.

In contrast to the extended credit model, the chunking model solved the *natural market problem* successfully in a wide range of client distributions (Fig. 4A, purple lines). To solve this task, the chunking model only needs to create the chunk *VR*, which in turn imposes a preference for the visitor, as *VR* is always associated with two units of food reward. The time of creating the *VR* chunk may vary according to the stochastic order of the trials experienced by each individual (see examples in Fig 4B). But on average, as the simulation advances, the chances of a cleaner using the chunking model to create the *VR* chunk and thus to choose a visitor increases (Fig. 4A, purple lines). Figure 4C depicts an example of the internal representation of the chunking model at the end of a simulation of the *natural market problem*. Note that the chunking model creates chunks regardless of the reward, and depending only on the statistics of cue occurrence. Thus, it may also create chunks containing the cue for an empty arena (*X*), or for other various combinations (e.g. *XR, XV, RR, VV*, etc.). In most cases these chunks do not influence the cleaner’s decisions as they represent states that require no choice (see Fig. 2). However, as we shall see below, in the natural setting there is also a risk of creating the *RV* chunk (rather than *VR* chunk), which can bias the cleaner’s decision, implying that chunking should be limited to avoid over-chunking.

### The fine-tuning of chunking behaviour and the effect of ecological conditions

The behaviour of the chunking model is controlled by the chunking avoidance parameter *C*_*p*_ (Eq. 2). Large values of *C*_*p*_ prevent any chunking and the model is reduced back to the core model (which do not develop a preference towards the optimal choice). On the other side, too low values of *C*_*p*_ cause ‘over-chunking’. Therefore, the optimal value of *C*_*p*_ will depend on the ecological conditions: the overall client density, and the frequency of the different client types. If there are many clients per cleaner, cleaners will often be solicited. Therefore, a visitor may regularly appear right after a resident – not because the visitor waits for service, but because a new visitor client enters into the arena by chance. Thus, there is a risk that the misleading chunk *RV* might be created, as well as the beneficial chunk *VR*. The reason we view the *RV* chunk as misleading is that faced with a choice between a visitor and a resident, the cleaner can now consider both sequences of actions, *V* →*R* and *R* →*V*, and choose between them according to their expected values. Although the value of the chunk *VR, f(VR)*, would approach 2 and hence be higher than the value of the chunk *RV* (with *f(RV)* lower than 2), the decision rule allows some proportion of choosing the *RV* chunk (exploration), which result in serving the resident first. In other words, over-chunking reduces the strength of the preference for the optimal choice. The balance between under-chunking and over-chunking implies the existence of optimal *C*_*p*_ values (balancing between the need to create the *VR* chunk but not the *RV* chunk). Importantly, these optimal *C*_*p*_ values depend on two ecological conditions: the overall client density, and the frequency of the different client types, which determine how frequently the sequences *V* →*R* and *R* →*V* are likely to be encountered. The effect of these ecological conditions on *C*_*p*_ and on the success of solving the *natural market problem* is shown in Fig. 5. It can be seen that in some extreme ecological conditions (of high client densities) it would be difficult for a cleaner fish using the chunking model to solve the market problem with any *C*_*p*_ (Fig. 5A, black dots, and Fig. 5B, blue shades representing low preference for visitors), since empty spots are rare events and most choices of the chunk *RV* result in obtaining two units of food (from the resident and the subsequent served client from the next trial). Fortunately for the cleaners, solving the market problem under these high client density conditions is not important in nature as high client densities lead to near permanent demand for cleaning. Yet, in most simulated ecological conditions where solving the market problem is important, an optimal *C*_*p*_ value (Fig. 5A) that induced a preference towards a visitor (Fig. 5B) was found.

**Figure 5.**
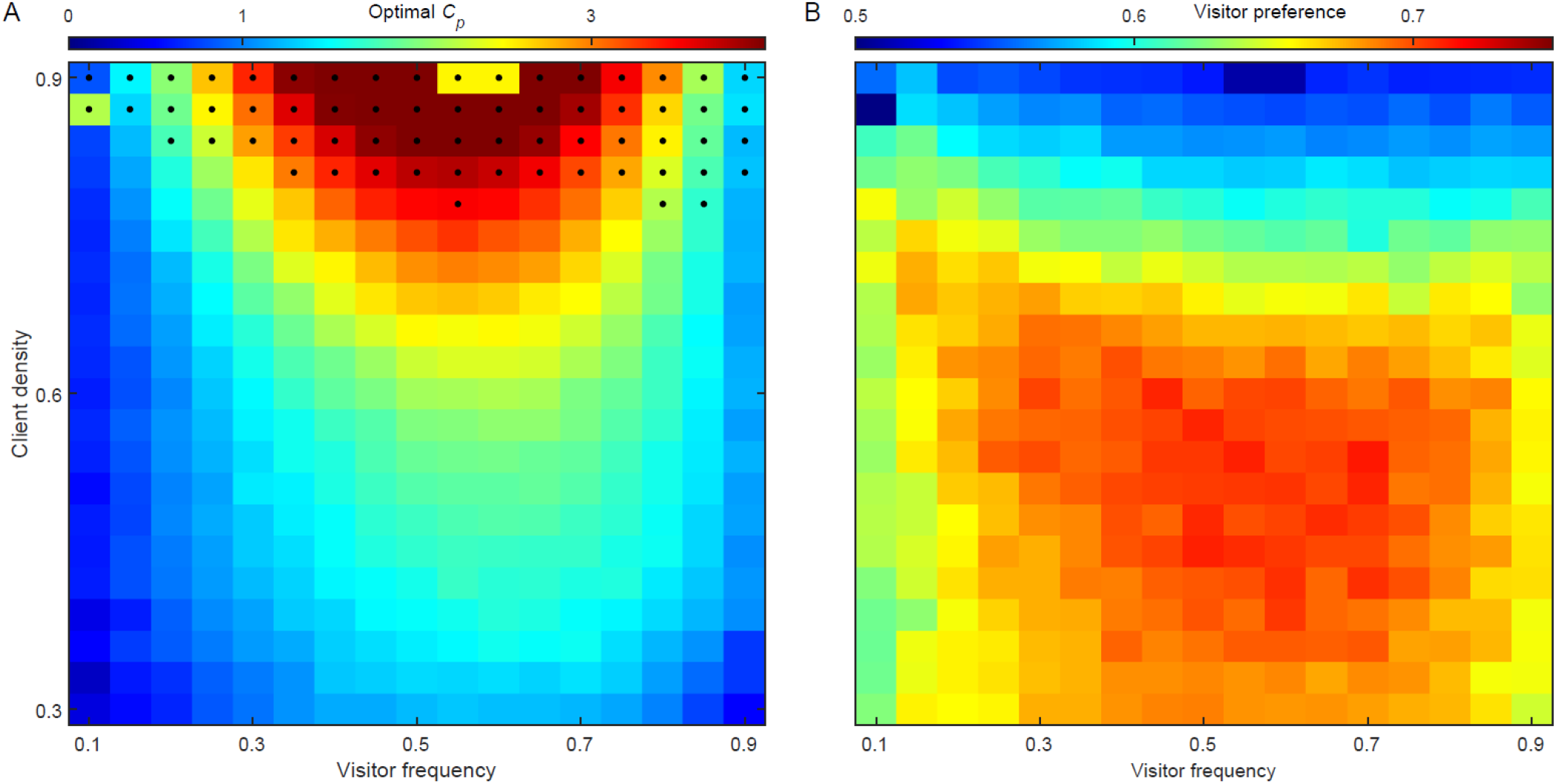
The link between ecological conditions, optimal *C*_*p*_, and the success of the chunking model in the *natural market problem*. **A)** Optimal *C*_*p*_ values (that provide the strongest preference towards a visitor), indicated by color, as a function of two ecological conditions: the visitor frequency,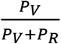 (the fraction of visitors out of all clients), and the overall client density, 1 −*P*_0_. The *C*_*p*_ values were estimated by running the simulations with 1000 values equally distributed between 0 and 5, fitting a Gaussian to the resulting visitor’s preferences, and finding its peak. Black dots depict conditions in which even the optimal *C*_*p*_ values resulted in a preference of less than 0.6 towards the visitor client. **B)** The preference (colour) towards the visitor client when the optimal *C*_*p*_ values are used in different ecological conditions.

To visualize the importance of the over-chunking problem, Figure S1 presents the frequency of appearances of each possible chunk (among 100 simulations) in four different ecological conditions, showing that when the chances of generating both the *VR* and *RV* chunks are similar (Fig. S1B), the preference towards visitors vanishes (compare with the relevant point of 0.5 visitor frequency and 0.9 client density in Fig. 5B).

Finally, our simulations show a significant positive correlation (linear regression: *R*^*2*^=0.78, *p*<0.001) between the frequency of simultaneous arrival of a visitor and a resident to the arena (hereafter: *r+v*; Fig. 6A) and the optimal *C*_*p*_ value (Fig. 6B). That is, when the combination of client density and visitors’ relative frequency increases the frequency of *r*+*v* pairs, a higher value of *C*_*p*_ should be used by the cleaners in order to increase the threshold of statistical significance allowing a chunk to be created. In contrast, when *r*+*v* pairs are rare, the probability of creating the misleading chunk *RV* is low so that lowering *C*_*p*_ is adaptive: it increases the likelihood of creating the beneficial chunk *VR* with almost no risk of creating the misleading chunk *RV*, which allows a strong preference for visitors to develop. Note, that when the frequency of *r*+*v* pairs is especially high (above 0.3; Fig. 6), there appears to be no *C*_*p*_ value that could balance between over-and under-chunking and the preference for visitors goes below 0.6 (only black dots appear at this range in Fig. 6B). More generally, a tendency to chunk too soon (e.g., *C*_*p*_ = 0.5) or too late (e.g., *C*_*p*_ = 2.5) resulted in poor performance under most combinations of client densities and visitor frequencies (Fig. S2).

**Figure 6.**
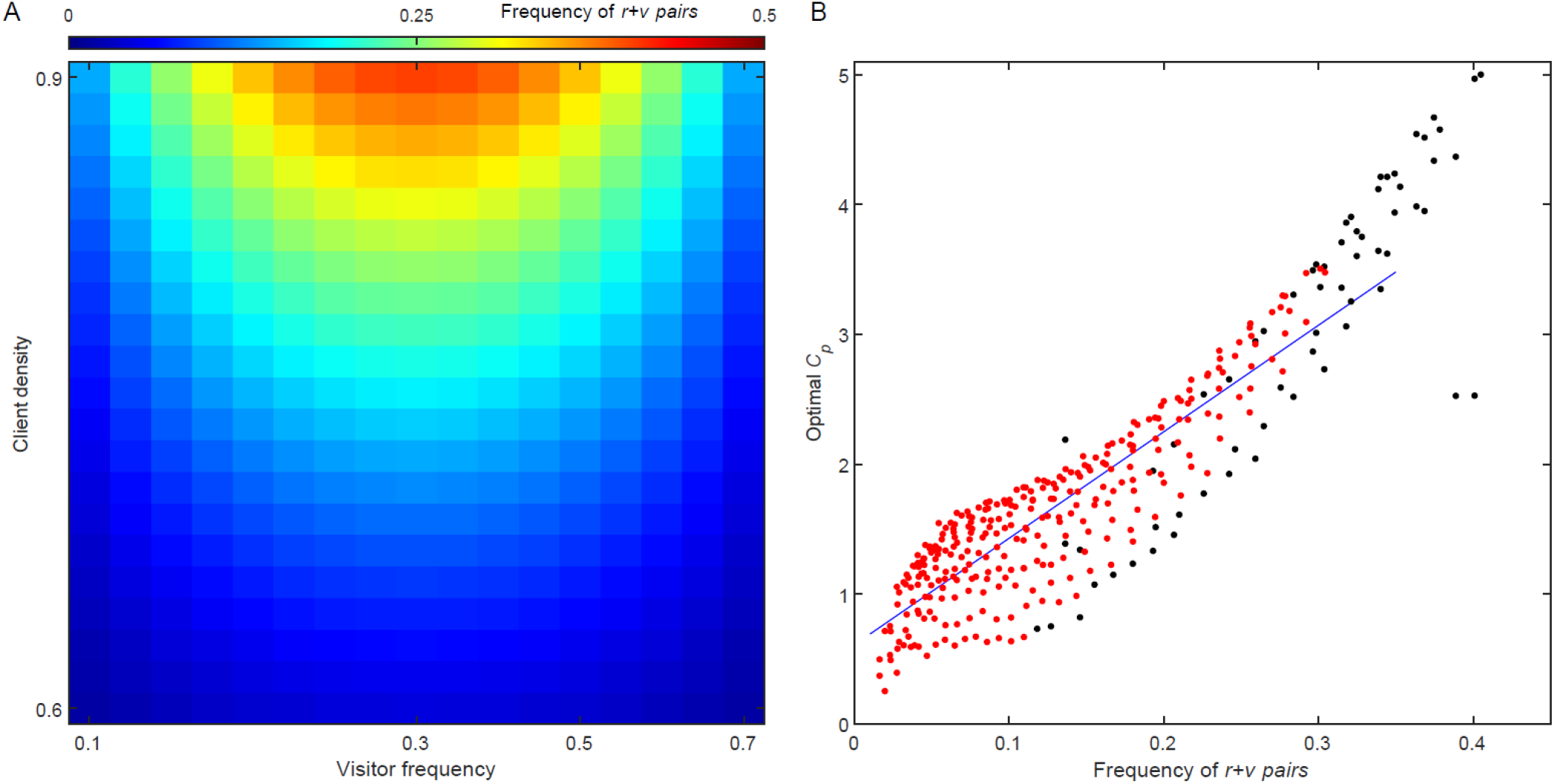
Correlation between Optimal *C*_*p*_ values and the frequency of resident and visitor pairs. **A)** The frequency of simultaneous appearance of resident and visitor (*r+v* pairs) in the arena, indicated by colour, out of all simulation trials (including empty and half empty trials) in the *natural market problem*. These are not stochastic values, but a feature of the simulated environment. **B)** The optimal *C*_*p*_ value (as in Fig 5A) as a function of the frequency of *r+v* pairs. Black dots – values obtained from simulations that achieved a preference towards a visitor lower than 0.6 (corresponding to the black dots in Fig. 5A). Blue line – linear regression of the optimal *C*_*p*_ values which achieved successful solutions (red dots; *R*^*2*^ = 0.78).

## Discussion

Chunking mechanisms are essential to represent structured data in the brain and have probably played a pivotal role in the evolution of cognition [28,30,33,41,42]. Yet, a possible challenge in the evolution of chunking is that incorrect chunking and over-chunking may lead to maladaptive behaviours and to cognitive impairments [66,67]. Indeed, the problem of under- or over-chunking arises whenever sensory input has to be chunked or segmented (reviewed in [33]). Normally, the problem is difficult to track because incoming data can be chunked in multiple ways and the number of possible chunks grows exponentially with the amount of data. This problem is well appreciated, for example, in the case of word segmentation during language learning in humans [32,68], or in the representation of behavioural sequences by animals [69]. Our analyses show that the market problem solved by cleaner fish in the wild offers a relatively simple model system to study the evolution of chunking. It is not only simple and tractable, but it involves a case where the function of chunking and its fitness consequences are well understood and are ecologically relevant, the adaptive and maladaptive chunks can be clearly identified (i.e., *VR* versus *RV*), and it can be studied experimentally and in relation to variable ecological conditions (e.g., [44,50,51]).

We implemented this approach by placing the same general problem of making a decision that doubles food intake in different sequential contexts that cleaners face in the wild. We show how solutions depend on increasingly complex learning rules. A simple two-choice task can be solved with basic reinforcement learning models such as the linear-operator or our equivalent core model. A more challenging task where doubling the amount of food is consistently due to consequences of an initial choice (i.e., the *laboratory market task*) requires an extended-credit learning model that picks up a consistent chain of events. Finally, if cleaners face diverse sequences of events, as in the *natural market problem*, relevant causal chains of subunits that lead to doubling the food intake must be identified and chunked so that the animal can optimise food intake.

We also demonstrate that when facing diverse sequences of events, having the ability to chunk may not be sufficient. It is critical that the decision to create chunks, captured by the chunking parameter *C*_*p*_, be adjusted to ecological conditions. Moreover, our simulations also show that under some extreme conditions, even the optimal chunking parameter may not be sufficient for developing a preference for the ephemeral reward. In the cleaners’ market problem, it happens when the probability of encountering the sequences of the useful and misleading chunks, *VR* and *RV*, respectively, is so similar that no chunking parameter can allow the creation of *VR* while preventing the creation of *RV*. As mentioned earlier, in the case of the cleaner fish, this may not be a problem because it happens under conditions of high client densities where preferring the ephemeral reward (i.e. visitors) is not necessary. It is yet to be studied how common are such conditions in other problems animals face in nature, and to what extent using the right chunking parameter is sufficient for successfully balancing the trade-off between under- and over-chunking.

Demonstrating the trade-off between adaptive chunking and over-chunking yields a new perspective on the cognitive basis of cleaner fish ‘cleverness’ in their choices of clients. Solving the natural market problem does not represent an “all or none” cognitive ability but rather the ability to correctly adjust a more basic cognitive ability, which is the ability to create chunks. As it stands, many animals are capable of creating chunks and configurations in their memory representation (see Introduction), but only those applying the chunking parameters suitable to the required conditions will solve the natural market problem. The trade-off between chunking and over-chunking may also explain why chunking (and configurational learning) takes time and may thus be viewed as difficult. Our model suggests that there is nothing really difficult in creating chunks quickly but that the process of chunking evolved to be slow in order to prevent over-chunking. Note that the idea that learning may evolve to be slow as a result of a trade-off is not new. It is implied in the optimization of learning rate parameters to balance between exploration and exploitation in reinforcement learning models [70,71], and was also suggested as a way to minimize recognition errors [72,73].

### The mechanism of chunking

Our chunking model specifies the statistical conditions required for the formation of chunks and describes how chunks are represented in the network (Eqs. 2 and 3; Fig. 1B). Yet, it does not explain how chunks are actually created. In other words, it does not explain how it happens that under the conditions specified by Equations 2 and 3, a chunk in the network suddenly appears. While the neuronal coding of such information is still poorly understood [29], a fairly explicit implementation of the process of chunk formation using neuronal-like processes may be possible. We can think of the required number of co-occurrences of *V* and *R* that is represented by the left side of Eq. 2 as the weight of their associative strength. Accordingly, a chunk representing the sequence *VR* is created when the weight of the edge leading from *V* to *R* passes a certain threshold. The formation of a chunk may be a result of another node in the network that receives signals from both neuronal units (or more precisely, from *R* soon after *V*), and thus increases in weight and becomes the “chunk node” representing the repeated occurrences of the sequence *VR* (as in Fig. 1B). The threshold weight required for the creation of a chunk can thus act as the chunking parameter *C*_*p*_ in our model and be optimized in line with Eq. 2.

In our model, that was kept as simple as possible, we assumed that weight increases by one unit per observation and does not decay over time. Realistically, however, different combinations of weight adjustment rates determine the timing crossing the threshold for chunk formation. For example, slow increase in weight with a relatively fast decay require frequent co-occurrences in order to reach the threshold, creating a test for the chunk’s statistical significance [19,33,41]. Thus, the chunking parameter in our model can be implemented by several mechanisms. We can hence view this parameter (or parameters) more generally as those effecting the tendency to form chunks (or the tendency to use configurational rather than elemental learning).

The optimization of the chunking parameters to ecological conditions may occur over generations through selection acting directly on parameter values, or instead (or in addition) cleaners may have evolved phenotypic plasticity with respect to the chunking parameter. For example, a rule instructing the cleaner to vary (loosen) the chunking parameters (i.e., explore) when in poor conditions and stop altering it (fasten) when in good conditions (i.e., exploit) may bring the chunking parameters to get fixated around the values associated with best performance. Another possibility is that cases where a visitor is leaving without waiting are experienced by the cleaner as aversive (a loss of a meal) and the aversive saliency of such events has evolved to reduce the chunking threshold (which increases the likelihood of chunking when solving the market problem is indeed necessary).

### Implications of our results on the interpretation of empirical studies

A major insight from our model in comparison to Quiñones et al. [62] is that animals only need the ability to detect chains of events (rather than chunking) in order to solve the laboratory market problem. Accordingly, it is not at all clear that differences between species in performance in the laboratory market task are due to different chunking abilities or different values of chunking parameters. It is hence important to use a more complex design of the market task (which resemble the natural setting for which chunking is necessary) on species that have solved some form of the laboratory task, i.e. cleaner fish, African grey parrots and capuchin monkeys [54,59]. Truskanov et al. [74] designed such a task, exposing cleaner fish to 50% of presentations of visitor and resident (*r+v*) plates as well as to 25% *r+r* and 25% *v+v* presentations. While a few cleaners solved this task, overall performance tended to be lower than in the standard laboratory market task. Applying our learning models to this non-standard (complex) market task showed that the extended credit model yields at best a slight preference for visitors, while the chunking model yields high performance (see Supplementary Information and Fig. S3). The study by Truskanov et al. thus yields experimental evidence that (some) cleaner fish can chunk. The task could also be adapted to test whether imposing “early commitment” that helped pigeons in solving the standard laboratory market problem [60] can also help to solve the natural problem, for which chunking ability is needed. Alternatively, “early commitment” can only help in extending the credit given to the initial choice (to the second reward as well as to the first one), which can solve the laboratory market problem but not the natural one.

Based on our model and simulations, there are currently multiple ways to explain the documented intraspecific variation in cleaner fish performance in both the standard and the complex laboratory market tasks [46,48,51,74]. First, variation in the laboratory market task may be related to whether individuals solve the problem by chunking or by chaining (extended credit) mechanisms, and to individual variation in the fine-tuning of the parameters of each mechanism. Second, assuming that cleaners use chunking to solve the tasks, variation in their performance may be attributed to some limitations or time lags in optimizing the chunking parameters to current conditions in the field, or to the specific conditions in the lab. Such limitations and time lags are expected for both genetic and phenotypically plastic adjustments because in the cleaners’ natural habitat, client densities and visitor frequencies can vary greatly across years and microhabitats [48,51], causing both inter- and intra-individual variation within individual lifetimes.

Importantly, these interpretations make related assumptions amenable for future testing. For example, that fast-solving cleaners use chunking even in the laboratory market task even though chaining would suffice, and that cleaners apply their field experience and developed *C*_*p*_ value to the lab task. Some empirical results are already in line with the second assumption. First, the best predictor of high cleaner performance in the laboratory task is high cleaner fish density [46], which in terms of our model implies low client density (per individual cleaner) and therefore low optimal *C*_*p*_ that promotes faster chunking (see Fig. 5A and Fig. 6). Second, individuals with relatively larger forebrains are more likely to be found in areas where they frequently face the market problem; [75]. Third, on a local scale, individuals with relatively larger forebrains performed according to what appears to be the locally best strategy: to solve the task if living in a high cleaner density area, and to fail the task if living in a low-cleaner density area [76]. In terms of our model, such high and low cleaner densities correspond to relatively low and high client densities that favour low and high *C*_*p*_ values, respectively (see Fig. 5A). Bringing such cleaners to lab implies that those who developed low *C*_*p*_ in their natural habitat are more likely to pass the test than those who developed high *C*_*p*_, which may explain Triki et al.’s results [76].

### Conclusions and implications for the study of advanced cognitive abilities

The cleaner fish ability to solve the market problem has presumably evolved on the background of its unique ecology and may be rightfully viewed as a surprisingly advanced cognitive ability for a (small brain) fish. However, by modeling the learning mechanisms required for this remarkable ability, we tried to put the cleaner fish story within the broader context of cognitive evolution, viewing it as a potential model for the evolution of chunking mechanisms. While the importance of chunking is usually considered within cognitive systems that are already highly advanced, the simple setting of the market problem allowed us to explicitly analyze the process of chunk formation, elucidating the trade-off between creating useful and misleading chunks, and demonstrating the importance of adjusting the chunking parameters to ecological conditions. We hope that the approach taken here could eventually be applied in the study of other cognitive abilities, identifying the learning mechanisms and the fine-tuning of their parameters required for their success, and mapping them not only along phylogenetic trees but also along evolutionary axes of explicit incremental changes in learning and cognitive mechanisms.

## Competing interests

The other authors declare that there are no competing interests.

## Supporting Information

### Laboratory complex market problem

Following the experiments conducted by Truskanov et al. [74] we simulated another environment of the market problem: the *lab complex market problem*. In this environment the cleaner faces a visitor-resident combination in 0.5 of the feeding trials, a resident-resident combination in 0.25 of the feeding trials, and a visitor-visitor combination in 0.25 of the feeding trials. As in the standard *laboratory market problem*, each feeding trial is followed by an empty trial. In the *lab complex market problem*, the extended-credit model generates only weak preference towards the visitor (Fig. S3, yellow line), yet it still did better than the core model or the linear operator that choose the clients with equal probabilities (Fig. S3, orange and blue lines). The reason for this minor preference is that in the *lab complex market problem* serving a second client after serving a visitor is still more frequent than serving a second client after serving a resident (even before any preference has been developed). This is because serving a visitor is followed by an empty trial only in the case of two visitors being presented (one is served and the other leaves), while serving a resident is followed by an empty trial in all combinations and choices except for a resident followed by another resident (in the visitor-resident choice, choosing a visitor means the resident is served last and choosing a resident also makes it last as the visitor leaves, while in the resident-resident case one is being served first and then the second resident is followed by an empty trial). Thus, the extended-credit model would assign somewhat higher value to the visitor and, subsequently, can develop some preference towards the visitor according to its decision rule.

**Figure S1.**
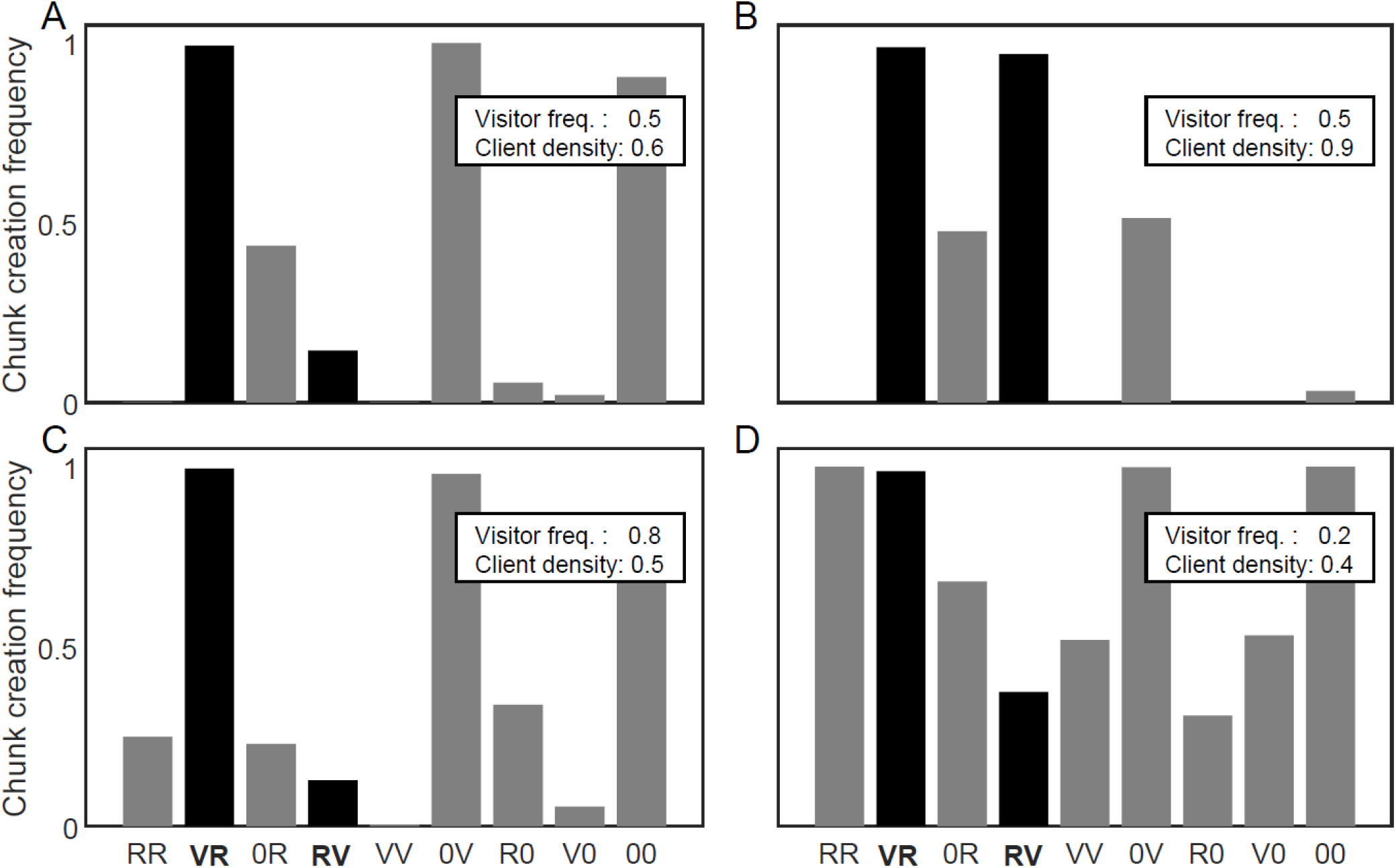
Creation of different chunks as part of the internal representation of the model in different ecological conditions. Four examples of ecological conditions are presented: **A)** visitor frequency of 0.5 and client density of 0.6, **B)** visitor frequency of 0.5 and client density of 0.9, **C)** visitor frequency of 0.8 and client density of 0.5, and **D)** visitor frequency of 0.2 and client density of 0.4. 1000 simulations were executed using the optimal *C*_*p*_ value for each condition (see Fig. 5A). The frequency of simulations, out of all simulations, in which the chunk was created by the end of the simulation, is presented for each chunk. Black bars – chunks which are relevant for the decision process; grey bars – chunks which are irrelevant for the decision.

**Figure S2.**
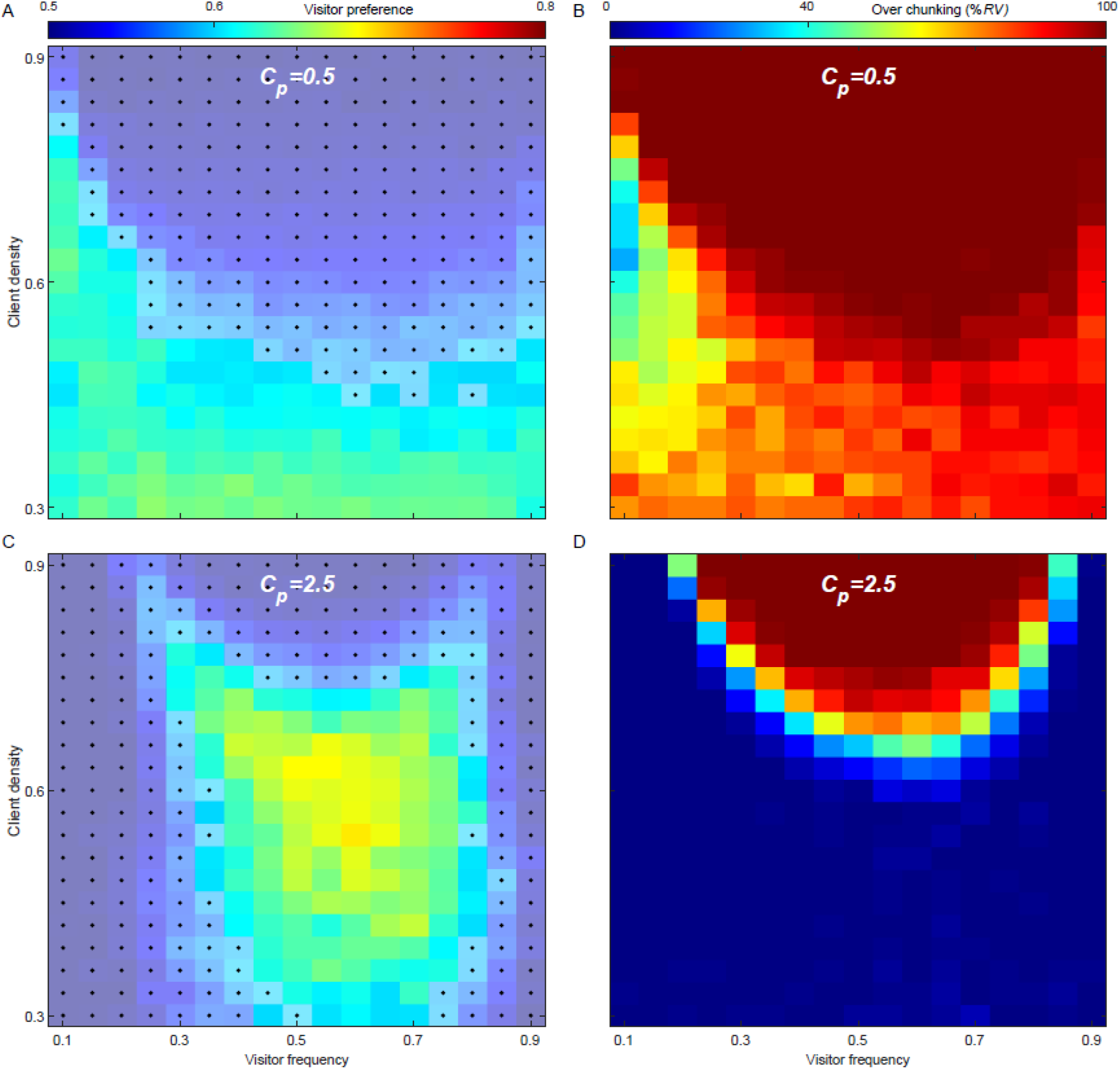
The link between ecological conditions, the success of the chunking model in the *natural market problem* using high and low *C*_*p*_ values, and over-chunking. **A)** The preference (colour) towards the visitor client when the *C*_*p*_ = 0.5 (low value) is used in different ecological conditions: the visitor frequency,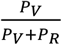 (the fraction of visitors out of all clients), and the overall client density, 1 − *P*_0_. The preference at each point is the mean of 100 simulations. Light colors with black dots depict conditions in which the preference towards the visitor client is less than 0.6. **B)** The percentage of simulations which ended up with the model generating the maladaptive *RV* chunk (over-chunking), when *C*_*p*_ = 0.5. Computed using 100 simulations for each point. **C)** The preference towards the visitor client when the *C*_*p*_ = 2.5 (high value) is used in different ecological conditions. **D)** The percentage of simulations which ended up with the model generating the maladaptive *RV* chunk (over-chunking), when *C*_*p*_ = 2.5. Note that low *C*_*p*_ and high *C*_*p*_ are beneficial under different conditions. Over-chunking is the cause of failure in the low *C*_*p*_ case. On the other hand, in the high *C*_*p*_ case, over-chunking is responsible to failures only in some conditions (high client density), but under-chunking fails the model in other conditions (low visitor frequency).

**Figure S3.**
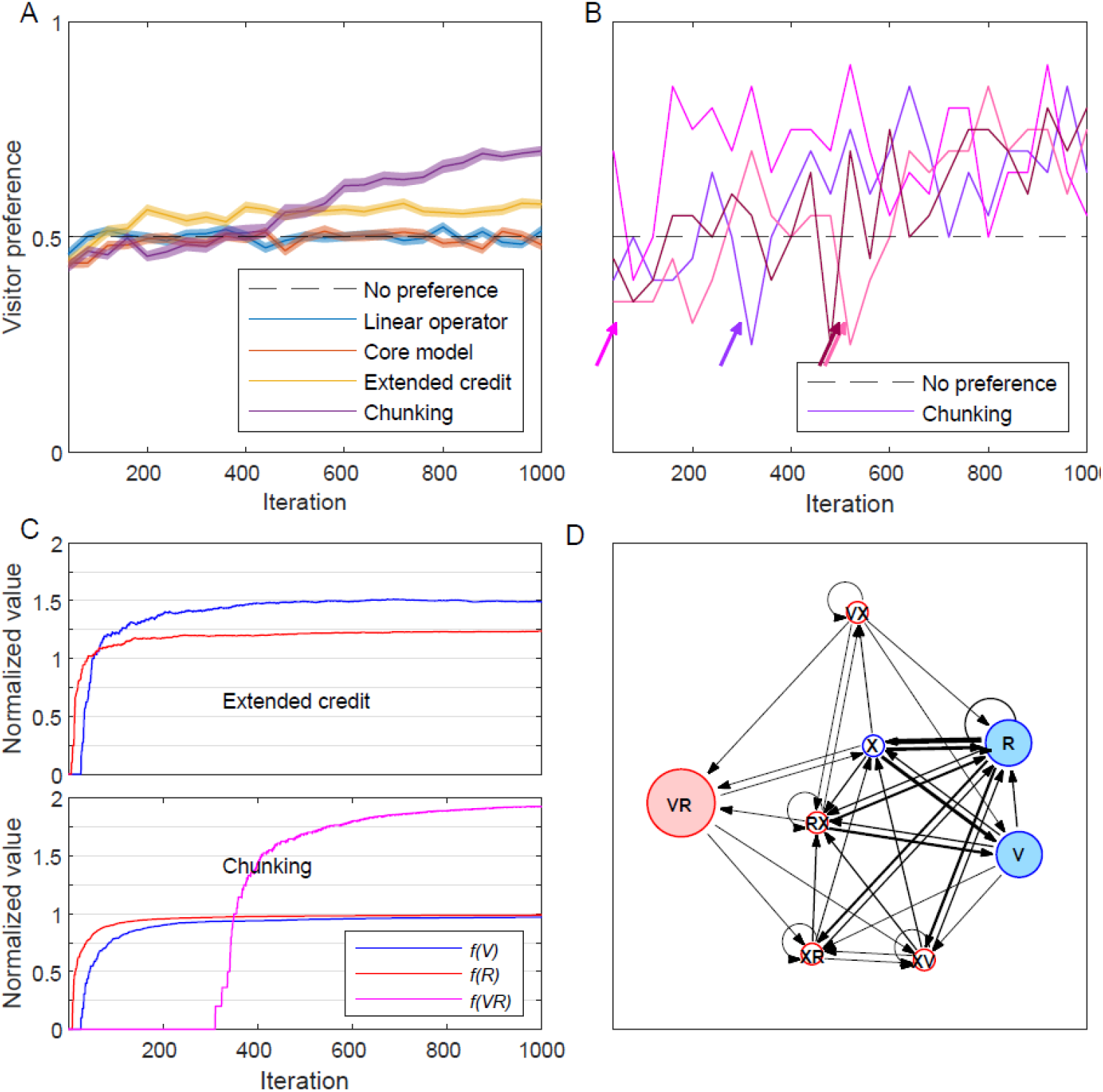
Simulating the *lab complex market problem*. **A)** Four types of learners are compared in the *lab complex market problem*: blue – A linear operator learner (α=0.1; see text); orange – the core model; yellow – the extended credit model; purple – the chunking model (with *C*_*p*_=2); black dashed-line – the expected choices with no preference (0.5). The preference towards a visitor client, measured as the proportion of choosing a visitor out of all visitor-resident encounters, is plotted as a function of time (iterations), in bins of 40 trials. Simulations are of 1000 feeding trials (with an empty trial after each feeding trial). The plots depict the mean of 100 simulations for each learner (shades – standard error of the mean) **B)** Four simulations of the chunking model. Note how the preference towards a visitor sharply increases after the creation of the *VR* chunk (depicted with an arrow for each simulation). **C)** The value of the different cues as perceived by the extended credit model (top) and the chunking model (bottom): blue – *V*; red – *R*; magenta – *VR*; in a single simulation. Note, how the extended credit model (top) converges towards a value of ∼1.5 for *V* and 1.25 for *R*, giving rise to a slight preference (∼0.6) towards a visitor (indicated by the yellow line in **A**; see text for discussion). **D)** The internal representation of the chunking model at the end of the simulation presented in (**C**, bottom). Blue – basic (initial) elements, red – chunk elements, filled nodes – the relevant elements for the decision process. The size of the circle is relative to the value (association with food reward) of the element. The width of the directed edges (black arrows) represents the relative frequency of the transitions between states (normalized *W*).

### Supplementary file

SimuFish.m – A Matlab function for running a simulation of the model in the cleaner fish market problem. See documentation inside.

## Notes

### Competing Interest Statement

The authors have declared no competing interest.

